# A cortical circuit for audio-visual predictions

**DOI:** 10.1101/2020.11.15.383471

**Authors:** Aleena R. Garner, Georg B. Keller

## Abstract

Learned associations between stimuli in different sensory modalities can shape the way we perceive these stimuli (Mcgurk and Macdonald, 1976). During audio-visual associative learning, auditory cortex is thought to underlie multi-modal plasticity in visual cortex (McIntosh et al., 1998; Mishra et al., 2007; Zangenehpour and Zatorre, 2010). However, it is not well understood how processing in visual cortex is altered by an auditory stimulus that is predictive of a visual stimulus and what the mechanisms are that mediate such experience-dependent, audio-visual associations in sensory cortex. Here we describe a neural mechanism by which an auditory input can shape visual representations of behaviorally relevant stimuli through direct interactions between auditory and visual cortices. We show that the association of an auditory stimulus with a visual stimulus in a behaviorally relevant context leads to an experience-dependent suppression of visual responses in primary visual cortex (V1). Auditory cortex axons carry a mixture of auditory and retinotopically-matched visual input to V1, and optogenetic stimulation of these axons selectively suppresses V1 neurons responsive to the associated visual stimulus after, but not before, learning. Our results suggest that cross-modal associations can be stored in long-range cortical connections and that with learning these cross-modal connections function to suppress the responses to predictable input.

---

Audio-visual interactions occur at many levels of cerebral processing from sub-cortical to high-level association cortices (Cappe et al., 2011), and thus the existence of direct connections between low-level sensory cortices (Falchier et al., 2002; Ibrahim et al., 2016; Iurilli et al., 2012) remains a curious enigma. Auditory input is known to influence neural activity in V1 (Fishman and Michael, 1973; Morrell, 1972; Murray et al., 2016; Petro et al., 2017). Some of these cross-modal responses are thought to be driven by direct projections from auditory cortex that target local inhibitory circuits in V1 (Ibrahim et al., 2016). While the computational role of these interactions remains unclear, we hypothesized that long-range cortical connections are shaped by experience and function to store memories of stimulus associations. Using an audio-visual associative conditioning paradigm, we investigated how cross-modal interactions shape neural responses in V1 over the course of learning. Mice explored a virtual environment in which they were exposed to sequentially paired presentations of auditory and visual stimuli. A virtual environment was used to enable simultaneous head-fixed optical physiology and experimental control of both visual and auditory input. Over the course of five conditioning sessions (approximately 45 minutes each on 5 consecutive days), mice were presented with pairings of a 1 second auditory cue (A) followed by a 1 second visual stimulus (V) (**Fig. 1a, b**). For each mouse, two pairs of an auditory cue and a visual stimulus were presented throughout conditioning (A_a_V_a_ and A_b_V_b_). The specific identity of stimuli used were counterbalanced across mice. To quantify the responses to the visual stimuli without a preceding auditory cue we occasionally presented the normally-cued visual stimuli alone (V_a_ and V_b_) and also presented a control visual stimulus (V_c_) that was never paired with an auditory cue. On day 5 of the conditioning paradigm, on a subset of trials, we additionally probed responses to an auditory cue and visual stimulus pairing the mouse had previously not experienced (A_b_V_a_). All presentations were randomized with an inter-stimulus interval between 4 and 12 seconds (see Methods).

**Figure 1.**
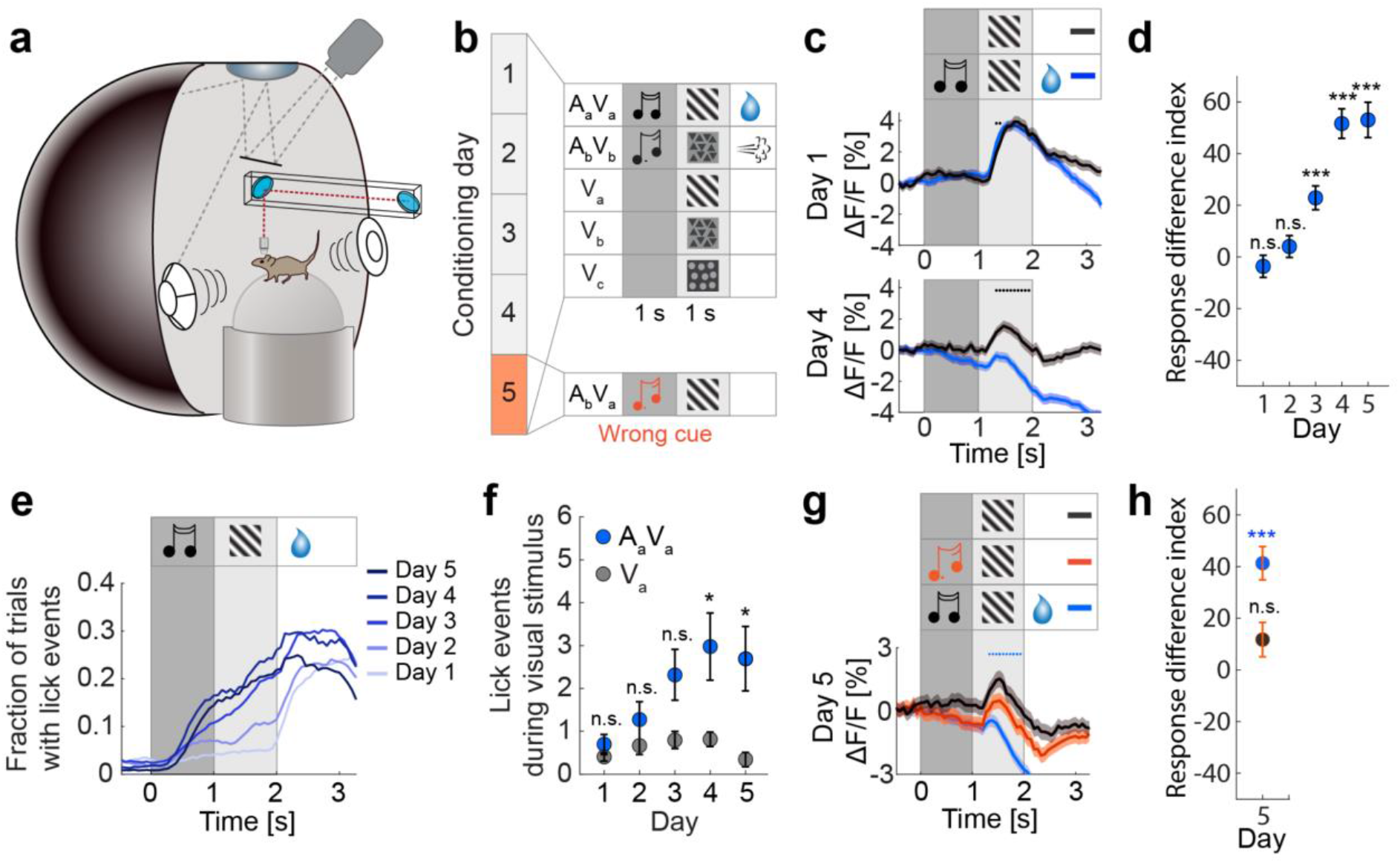
Visual responses are suppressed by an associated auditory cue. **(a)** Schematic representation of the virtual reality setup. **(b)** Experimental paradigm. Over the course of 5 conditioning days, mice were exposed to auditory-cued visual stimuli (A_a_V_a_, A_b_V_b_) that were reinforced, to the visual stimuli alone (V_a_, V_b_) with no reinforcement, and to a control visual stimulus (V_c_) that was never paired with an auditory stimulus or reinforced. On day 5, mice were additionally exposed to a previously unexperienced audio-visual stimulus pair (A_b_V_a_). **(c)** Average population responses of L2/3 V1 neurons for cued (A_a_V_a_, blue) and un-cued (V_a_, gray) visual stimulus presentations on day 1 (top) and day 4 (bottom) of conditioning. For **c, d**, and **g** day 1 - 4: n = 1548 neurons from 10 mice; day 5: n = 1341 neurons from 9 mice. Black dots indicate traces are different during visual stimulation (p<0.05, paired two-sided t-test, see Methods for detailed calculation). Here, and in the following figures, the dark gray bar indicates auditory stimulus presentation, and the light gray bar indicates visual stimulus presentation. **(d)** Quantification of the difference in response for each conditioning day (Response difference index) during the auditory-cued and un-cued visual stimulus presentations, normalized by the mean response during the un-cued visual stimulus on day 1 (V_a_-A_a_V_a_)/ mean(V_a_). Asterisks indicate comparison to 0 difference using a two-sided rank-sum test. Here and in subsequent panels *: p <0.05, **: p <0.01, ***: p <0.001; for all statistical analyses and exact p values see **Table 1**. **(e)** Anticipatory licking increases with conditioning day for A_a_V_a_. Traces indicate mean fraction of trials with lick events. For **e** and **f**, day 1 - 4: n = 10 mice and day 5: n = 9 mice. **(f)** Anticipatory licking for A_a_V_a_ (blue) and V_a_ (gray) with conditioning as quantified by lick events during visual stimulus presentation. Mean ± SEM across mice. Asterisks indicate comparison between A_a_V_a_ and V_a_ trials using a two-sided rank-sum test. **(g)** Mean population responses on day 5 on which a subset of trials consisted of previously unpaired stimuli (A_b_V_a_). The response during A_b_V_a_ (orange) was different from the response during A_a_V_a_ (blue) but not from the response during V_a_ (gray). Blue dots indicate A_b_V_a_ and A_a_V_a_ curves are different (see Methods). **(h)** Quantification of the difference in responses in **g** (Response difference index). The response during the visual stimulus of condition A_b_V_a_ is greater than that during condition A_a_V_a_ (blue with orange), but not different from the response during V_a_ (gray with orange). Comparisons were made using a two-sided rank-sum test. For neural activity plots, traces or filled circles indicate the mean and shading or error bars indicate SEM across neurons.

The behavioral relevance of visual stimuli is known to influence the dynamics of neural responses in V1 in paradigms in which the animal is exposed to the same stimuli over the course of days (Henschke et al., 2020; Keller et al., 2017; Poort et al., 2015). To test the influence of the behavioral relevance of the paired stimuli, we performed two variants of the conditioning paradigm in two groups of mice, one in which the paired stimuli were followed by appetitive or aversive reinforcements, and one in which the paired stimuli were not reinforced. In the reinforced variant, A_a_V_a_ was followed by a water reward and A_b_V_b_ by a mild air puff to the neck. To monitor neural activity, three weeks prior to the conditioning experiments, we injected an adeno-associated viral (AAV) vector expressing a genetically encoded calcium indicator (AAV2/1-EF1α-GCaMP6f) in right monocular V1. Throughout conditioning, mice were head-fixed on a spherical treadmill and free to locomote. Rotation of the treadmill was coupled to movement in a virtual tunnel displayed on a toroidal screen surrounding the mouse. The precise location of V1 in retinotopic coordinates was measured for all mice using optical imaging of intrinsic signals (**Extended Data Fig. 1a**). We recorded neural activity in layer 2/3 (L2/3) of V1 using two-photon calcium imaging. Visual stimuli were presented bilaterally in visual space matched to the retinotopic location of the two-photon imaging region. Auditory stimuli were presented through a speaker pair located symmetrically on either side of the mouse.

## Visual responses are suppressed by an associated auditory cue

To first assess the effect of repeated exposure to a visual stimulus over the course of conditioning, we examined population responses to V_c_, which was never paired with an auditory cue or reinforced, and found a general decrease in responsiveness across days (**Extended Data Fig. 1b**). To test whether an audio-visual association affected whether V1 responded differently to a visual stimulus, we compared the average population responses to the auditory cue and visual stimulus pair that was followed by a reward (A_a_V_a_), to that of the same visual stimulus (V_a_) presented alone. We found that on day 1 of conditioning, the two visual responses were similar (**Fig. 1c**). Analogous to V_c_, over the course of conditioning, the visual responses to both A_a_V_a_ and V_a_ decreased (**Extended Data Fig. 1c**). Interestingly, however, we found that the auditory cue preceding the paired visual stimulus resulted in an additional suppression of the visual response that increased with experience (**Figs. 1c, d, Extended Data Fig. 1c**). Furthermore, this suppression was most prominent for the auditory and visual stimuli followed by a water reward. For the audio-visual stimuli followed by an air puff (A_b_V_b_), we also observed a suppression of the visual response following the auditory cue, however this suppression developed already on day 1 and was weaker and more variable than in the rewarded condition (**Extended Data Figs. 1d, f**). Additionally, the auditory cue itself resulted in a slight increase in V1 activity initially and a slight decrease in activity later in conditioning (**Extended Data Fig. 1e**). In mice that underwent the same pairing paradigm without any reinforcements, visual responses were smaller on average (**Extended Data Fig. 1g**), and the auditory cue did not result in a consistent suppression of the visual response (**Extended Data Figs. 1g, i**). Similar to reinforced conditioning, the auditory cue itself initially resulted in a slight increase in activity, but unlike reinforced conditioning, this response did not change over time (**Extended Data Fig. 1h**). To investigate the mechanism of auditory-cue driven suppression of visual responses, we focused subsequent analyses on the stimuli that were reinforced with a water reward. In addition to the experience-dependent auditory-cue driven suppression, we also found that the visual responses to A_a_V_a_ and V_a_ decorrelated with experience (**Extended Data Fig. 2a**). Thus, learned audio-visual associations can change the way V1 represents visual stimuli depending on the behavioral relevance of the stimuli.

To test whether mice exhibited a conditioned behavioral response to the presentation of the audio-visual stimulus A_a_V_a_ preceding the water reward on a time-scale similar to that of the audio-visual suppression, we quantified licking responses in anticipation of the water reward. Over the course of conditioning days, mice successively increased the number of licks made before reward delivery during the presentation of the auditory-cued visual stimulus (**Fig. 1e**). Although the presentation of the visual stimulus in the absence of the auditory cue (V_a_) also resulted in occasional licking, this response was much weaker (**Fig. 1f**). To test whether auditory-cue driven suppression of visual responses was caused by a differential behavioral response during A_a_V_a_ and V_a_, we took advantage of the variability in licking behavior. While mice exhibited an increased licking response to the auditory-cued visual stimulus, they also exhibited licking in a subset of non-cued visual stimulus trials (day 1: 26.8% ± 5.3% of trials, day 4: 39.9% ± 8.4% of trials, mean ± SEM) and did not lick during a subset of the auditory-cued visual stimulus trials (day 1: 63.5% ± 9.2% of trials, day 4: 27.2% ± 9.4% of trials, mean ± SEM). We could thus compare the responses in trials with and without licking for both conditions separately (**Extended Data Fig. 2b**). On day 1, we found no response difference induced by licking. On day 4 licking also did not result in a reduction of the response to the visual stimulus when presented alone indicating that licking per se did not drive a suppression of the visual response. However, for the auditory-cued visual response, the suppression on day 4 was only present in trials in which the mouse exhibited anticipatory licks to the reward. Thus, once the association was established, the auditory cue only resulted in a suppression of the visual response when it was accompanied by a licking response. This suggests that mice must acknowledge presentation of the paired stimuli for the auditory cue to have a suppressive effect on the visual response. In parallel to the anticipatory licking responses, both auditory and visual stimuli induced a reduction in average locomotion speed (**Extended Data Fig. 2c**), which is known to modulate visual responses (Keller et al., 2012; Niell and Stryker, 2010; Saleem et al., 2013). However auditory-cue driven suppression was not explained by variance in locomotion, as it was still present in speed-matched trials (**Extended Data Figs. 2d, e**). Thus, differences in running speed cannot account for the observed experience-dependent suppression of the visual responses by the auditory cue.

To determine whether suppression of the visual response developed specifically for the auditory cue paired with the visual stimulus, we presented previously unpaired auditory cue and visual stimulus pairings in a subset of the trails on day 5 of conditioning (A_b_V_a_). We found that suppression of the visual response was specific to the auditory cue with which the visual stimulus had been paired. There was no suppression when the visual stimulus was preceded by a different auditory cue than the one with which it had been associated, and the response to the visual stimulus following a different auditory cue, A_b_V_a_, was not different from the response to the visual stimulus alone V_a_ (**Figs. 1g, h, Extended Data Fig. 2e**). In summary, we find that in a behaviorally relevant context, the association of an auditory cue with a visual stimulus results in a stimulus specific suppression of the visual response in L2/3 of V1.

## Auditory input to visual cortex is multi-modal and experience dependent

Visual and auditory cortices directly interact both anatomically and functionally (Bizley et al., 2007; Clavagnier et al., 2004; Falchier et al., 2002; Morrell, 1972; Shams et al., 2002; Zangenehpour and Zatorre, 2010) resulting in responses to visual and auditory stimuli in both regions (Clavagnier et al., 2004; Kayser et al., 2008; McIntosh et al., 1998). Auditory cortex projects directly to V1 in primates (Falchier et al., 2002; Majka et al., 2019) and rodents (Budinger and Scheich, 2009; Miller and Vogt, 1984), where it constitutes one of the densest inputs to V1, as quantified by rabies tracing in mice (Leinweber et al., 2017). To test whether direct projections from auditory cortex (AuC) to V1 could contribute to the auditory-cued suppression of visual responses, we repeated the conditioning experiments in a cohort of mice in which we functionally imaged AuC axons in V1. We injected an AAV2/1-EF1α-GCaMP6s vector in AuC to express GCaMP6s in AuC neurons and implanted an imaging window over ipsilateral V1 to perform two-photon imaging of superficial AuC projection axons in V1 (**Figs. 2a, b**). We confirmed in postmortem histological analysis that the vast majority of the neurons labelled were in AuC and that the few neurons retrogradely labelled in V1 could not account for the number of axons we recorded in V1 (**Extended Data Figs. 3a, b**).

**Figure 2.**
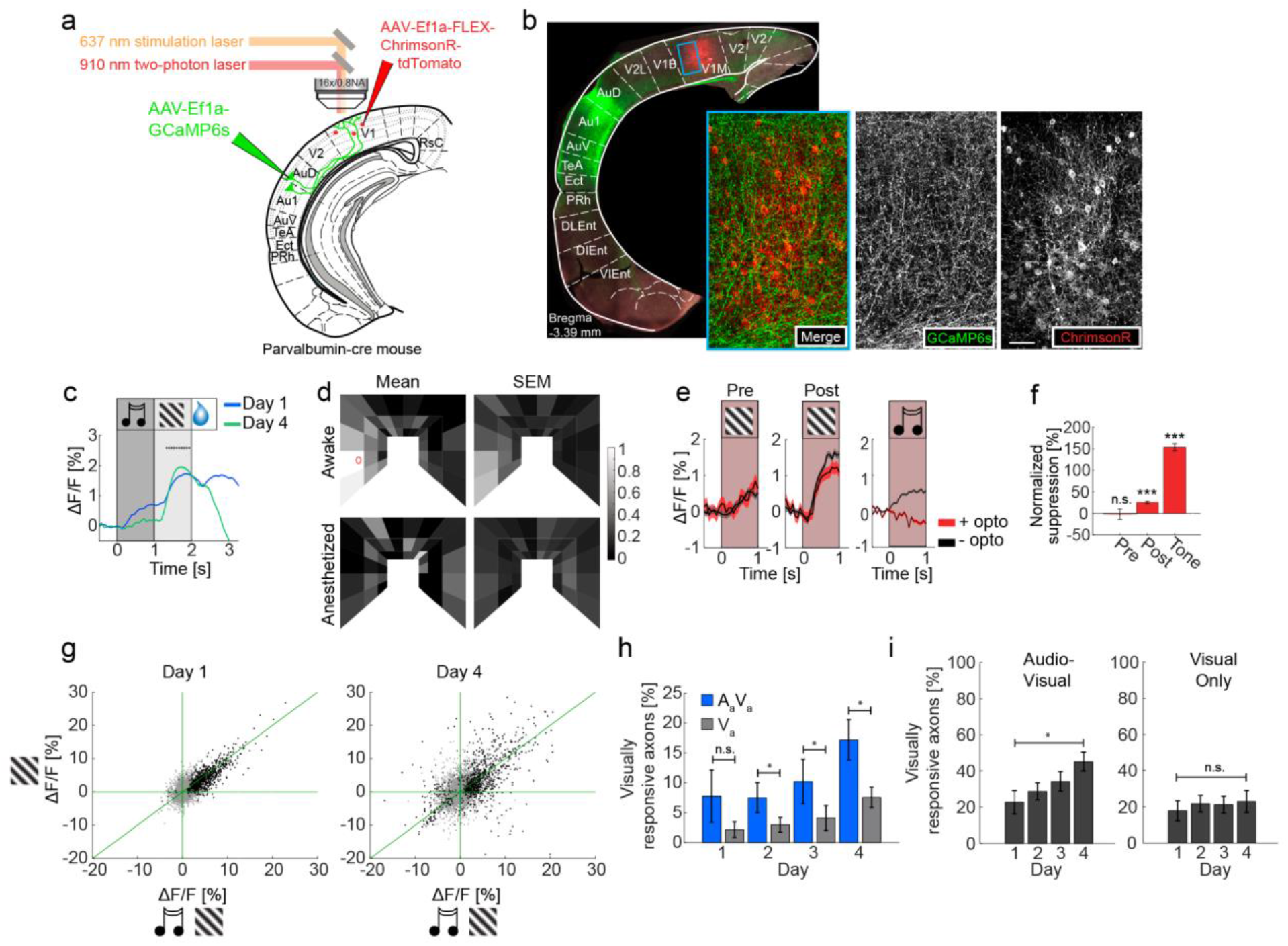
Auditory cortex (AuC) sends experience dependent audio-visual signals to V1. **(a)** Schematic of injection sites referenced to atlas (Paxinos and Franklin, 2013). GCaMP6s injection in AuC and ChrimsonR-tdTomato injection in V1. **(b)** Representative confocal histology image illustrating AuC axonal projections to V1 (green) and V1 PV neurons (red) at the approximate imaging location. Insets show region marked by blue box in V1. Scale bar indicates 50 µm. **(c)** AuC axons in V1 respond to the auditory cue and to the visual stimulus. Day 1: n = 21,076 axons from 20 mice, day 4: n = 19,486 axons from 19 mice. Also see **Extended Data Figs. 3c - e**. Black dots indicate traces are different during visual stimulation. (p<0.05, paired two-sided t-test, see Methods for detailed calculation). **(d)** Visual responses of AuC axons were mapped in a virtual corridor environment (see Methods). Visual responses of AuC projection axons were retinotopically matched to the imaging location in V1 in awake mice (top, 4305 axons in 7 mice). The red circle marks the average peak location of visual responses of V1 neurons recorded in the same anatomical location and the same stimulation setup (Zmarz and Keller, 2016). In anesthetized mice, visual responses were nearly absent (bottom, 991 axons in 5 mice). (Left column) mean responses plotted as a function of location in visual space in the virtual corridor. (Right column) corresponding SEM. Color scale is normalized to the peak response (1.1% ΔF/F). **(e)** Inhibiting V1 locally by optogenetic excitation of PV positive interneurons had no effect on visual responses before conditioning (left, 2927 axons in 7 mice), and a moderately suppressive effect after conditioning (middle, 3857 axons in 7 mice), but resulted in complete suppression of auditory responses (right, 4130 axons in 6 mice). Red bar indicates laser illumination. **(f)** Normalized suppression quantified as the difference between the response to the stimulus with and without optogenetic inhibition, normalized by the mean response to the stimulus without inhibition. Pre: n = 2927 axons from 7 mice, Post: n = 3857 axons from 7 mice, Tone: n = 4130 axons from 6 mice. Asterisks indicate comparison to 0% suppression using a two-sided rank-sum test. Here and in subsequent panels *: p <0.05, **: p <0.01, ***: p <0.001; for all statistical analyses and exact p values see **Table 1**. **(g)** Average visual response of each axon to A_a_V_a_ plotted against the visual response to V_a_ on day 1 (left) and day 4 (right). Black data points are axons with a significant response to either visual stimulus condition. For **g - i**, day 1: n = 5552 axons from 8 mice, day 2: n = 5097 axons from 8 mice, day 3: n = 5157 axons from 8 mice, and day 4: n = 4658 axons from 7 mice. **(h)** Fraction of visually responsive axons to A_a_V_a_ (blue) and V_a_ (gray) as a function of conditioning day. Comparisons were made using a paired two-sided t-test. **(i)** (Left) Fraction of visually responsive axons as a function of conditioning day in the audio-visual conditioning context. (Right) For the same mice and axons, in a visual only context, the fraction of visually responsive axons did not change from day 1 to day 4. Comparisons were made using an unpaired two-sided t-test. For neural activity plots, traces or bars indicate the mean and shading or error bars indicate SEM across axons. For quantification of visually responsive axons, bars indicate the mean and error bars indicate SEM across mice.

Recording the activity of AuC axons in V1, we found that early in conditioning these carried both an auditory response as well as a visual response (**Fig. 2c**). Interestingly, the visual responses were larger than the auditory responses and, different from responses in V1, increased slightly over the course of conditioning (**Fig. 2c, Extended Data Figs. 3c, d**). Conversely, the auditory responses in AuC axons, like the visual responses in V1, decreased across conditioning days (**Fig. 2c, Extended Data Fig. 3e**). Intrigued by the strength of the visual responses, we mapped the responses as a function of retinotopic location of the visual stimulus and found that they had receptive fields that matched the retinotopic location of the recording location in V1 (**Fig. 2d, top**). These visual responses were absent in anesthetized recordings (**Fig. 2d, bottom**) suggesting the visual responses may arise from cortico-cortical connections (Raz et al., 2014). Given that visual cortex also projects to auditory cortex (Bizley et al., 2007; Kayser et al., 2008), it is possible that the source of the visual responses in AuC axons is inherited from retinotopically matched V1 neurons. To test this, we examined AuC axon responses while silencing activity in V1 locally. We used a mouse line expressing Cre in parvalbumin (PV) positive interneurons (Hippenmeyer et al., 2005), and injected an AVV vector to express a Cre-dependent channelrhodopsin variant in V1 (AAV2/1-EF1α-DIO-ChrimsonR-tdTomato). We then quantified the effect of locally silencing V1 using optogenetic activation of PV interneurons while imaging the calcium responses in AuC axons (see Methods). Surprisingly, we found that the inhibition of V1 activity was effective in suppressing auditory evoked responses in the AuC axons but resulted in no suppression of visual responses before conditioning and only a small reduction after conditioning (**Figs. 2e, f**). The responsiveness of AuC projection axons to visual stimuli is consistent with previous work in awake mice showing visually responsive neurons in auditory cortex are predominantly found in layers 5 and 6 (Morrill and Hasenstaub, 2018), which send collaterals to cortical targets including V1 (Ibrahim et al., 2016). However, the role of visual responses in auditory cortex remains elusive. Our results demonstrate that AuC conveys a retinotopically matched visual signal to V1 largely independent of V1 activity. Such a signal could potentially function to inhibit the auditory-cued visual response in visual cortex. In order for AuC input to contribute to the experience-dependent suppression of auditory-cued visual responses, we would expect an experience-dependent change in the AuC axon responses over the course of conditioning. Congruently, we found that there was a decrease of similarity between axon visual responses to A_a_V_a_ and V_a_ between day 1 and day 4 of conditioning (**Fig. 2g**). In addition, we found that the fraction of visually responsive axons was greater when the visual stimulus followed the auditory cue (A_a_V_a_) than when presented alone (V_a_) (**Fig. 2h**). This result prompted us to examine differences in visual responsivity of AuC axons when mice were tasked with learning audio-visual associations compared to when they were similarly exposed only to visual stimuli. We therefore exposed the mice in our audio-visual conditioning context to a second context, over the same time course of conditioning, in which only visual stimuli were presented (see Methods). We found that while the overall fraction of visually responsive axons increased from day 1 to day 4 of conditioning in the audio-visual context (**Fig. 2i, left**), there was no difference in the fraction of visually responsive axons from day 1 to day 4 in the visual-only context (**Fig. 2i, right**). Thus, AuC input to V1 exhibits an experience-dependent modulation of the visual response by the auditory cue.

## Suppression of V1 by auditory input is stimulus and experience dependent

AuC input could functionally suppress the auditory-cued visual responses either by global suppression, independent of stimulus preference of neurons in V1, or by specific suppression of the neurons responsive to the visual stimulus paired with an auditory cue. In either case, given that the suppressive effects we observe in V1 are experience-dependent, we would also expect the suppressive action of AuC to be learned with experience. To test if the AuC input to V1 could function as either a global or a functionally specific suppressive input, we used an experimental paradigm to map the functional influence of the AuC input on functionally identified V1 neurons before and after conditioning. We injected a vector expressing a channelrhodopsin variant (AAV2/1-EF1α-ChrimsonR-tdTomato) in AuC and a vector expressing GCaMP6f (AAV2/1-EF1α-GCaMP6f) in V1 (**Fig. 3a**). This allowed us to functionally map the influence (FMI) of the AuC axon stimulation on neural responses of L2/3 V1 neurons. We used a 1 second pulse of a 637 nm laser to activate the ChrimsonR in the imaging region during two-photon imaging (see Methods). As the stimulation occurred optically coaxial with the two-photon imaging, the mouse’s eyes were shielded from stimulation light by the imaging cone. To control for a putative effect of the stimulation light directly driving a visual response, we also performed sham stimulations with a second light source diffusely illuminating the head of the mouse outside of the imaging cone. Stimulation of the AuC axons resulted in a variety of responses in V1 (**Fig. 3c**). In unconditioned mice, 37.7 ± 8.2 % of neurons were responsive to AuC axon stimulation and of these 48.4 ± 20.1 % were inhibited (n = 5 mice). In conditioned mice, 35.4 ± 7.0 neurons were responsive to AuC axon stimulation and of these 30.6 ± 11.1 % were inhibited (n = 10 mice). While we also observed a response to the sham stimulation, we found no correlation between the response to AuC axon stimulation and sham stimulation (**Extended Data Fig. 4a**), indicating that the response to the optogenetic stimulation of the AuC axons cannot be explained by a visual response. We then examined if an experience-dependent alteration of the connection from AuC to V1 existed in the form of a difference in the pattern of activation induced in V1 by the AuC stimulation before and after audio-visual experience. We tested this by functionally mapping the influence of AuC axon stimulation in the same L2/3 V1 neurons before and after conditioning (**Fig. 3b**). This allowed us to determine whether there was a relationship between the responses of a neuron to sensory stimulation (i.e. V_a_ and A_a_V_a_) and to the artificial activation of AuC projection axons and if there was an experience-dependent change in the influence of the AuC input on V1. The average V1 population response to artificial AuC activation remained similar before and after conditioning (**Extended Data Fig. 4b**). Plotting the response to the artificial AuC stimulation for every V1 neuron before conditioning against the response after conditioning revealed a variety of learning related changes that were larger than those expected simply from response variability to the stimulation on a trial-by-trial basis (**Fig. 3d, Extended Data Fig. 4c**). To examine whether the neurons with strong responses to either V_a_ or A_a_V_a_ were preferentially functionally influenced by the AuC axon stimulation, we color coded the response of each neuron to V_a_ and A_a_V_a_, early and late in conditioning on scatter plots of their responses to the AuC axon stimulation pre- and post-conditioning (**Fig. 3d**). We found that early in conditioning no correlation existed between responses to the visual stimulus and response to AuC axon stimulation. However, late in conditioning, neurons with the strongest excitatory responses to the visual stimulus tended to cluster in the lower-left quadrant of the scatter plot, meaning the neurons that were functionally inhibited by the stimulation of AuC axons showed the strongest responses to V_a_. Moreover, the visual responses of these neurons were reduced in the A_a_V_a_ condition. To quantify this tendency and examine the stimulus specificity of AuC axon stimulation effects, we split V1 neurons into those inhibited by and those excited by AuC axon stimulation and compared visual responses of these populations to the paired stimulus V_a_ or the control stimulus V_c_. Neurons with a decrease in fluorescence during AuC axon stimulation were classified as inhibited and those with an increase as excited. This definition allowed inclusion of all neurons in the analysis. While early in conditioning no difference existed between the mean visual responses of neurons either excited or inhibited by AuC axon stimulation, after conditioning, the neurons inhibited by AuC axon stimulation exhibited larger responses specifically to V_a_ (**Fig. 3e**). Importantly, the inhibited population showed the largest suppression of the visual response following the previously-paired auditory cue (**Extended Data Figs. 4d, e**) and the largest recovery of the visual response following the previously-unpaired auditory cue (**Extended Data Figs. 4f, g**). This is consistent with a specific targeting of the functional inhibition to neurons receiving the strongest drive from the visual stimulus that was paired with the auditory cue. Thus, experience reshapes the influence of the long-range cortical connection between AuC and V1 to suppress responses to visual stimuli the mouse learns to predict from auditory cues.

**Figure 3.**
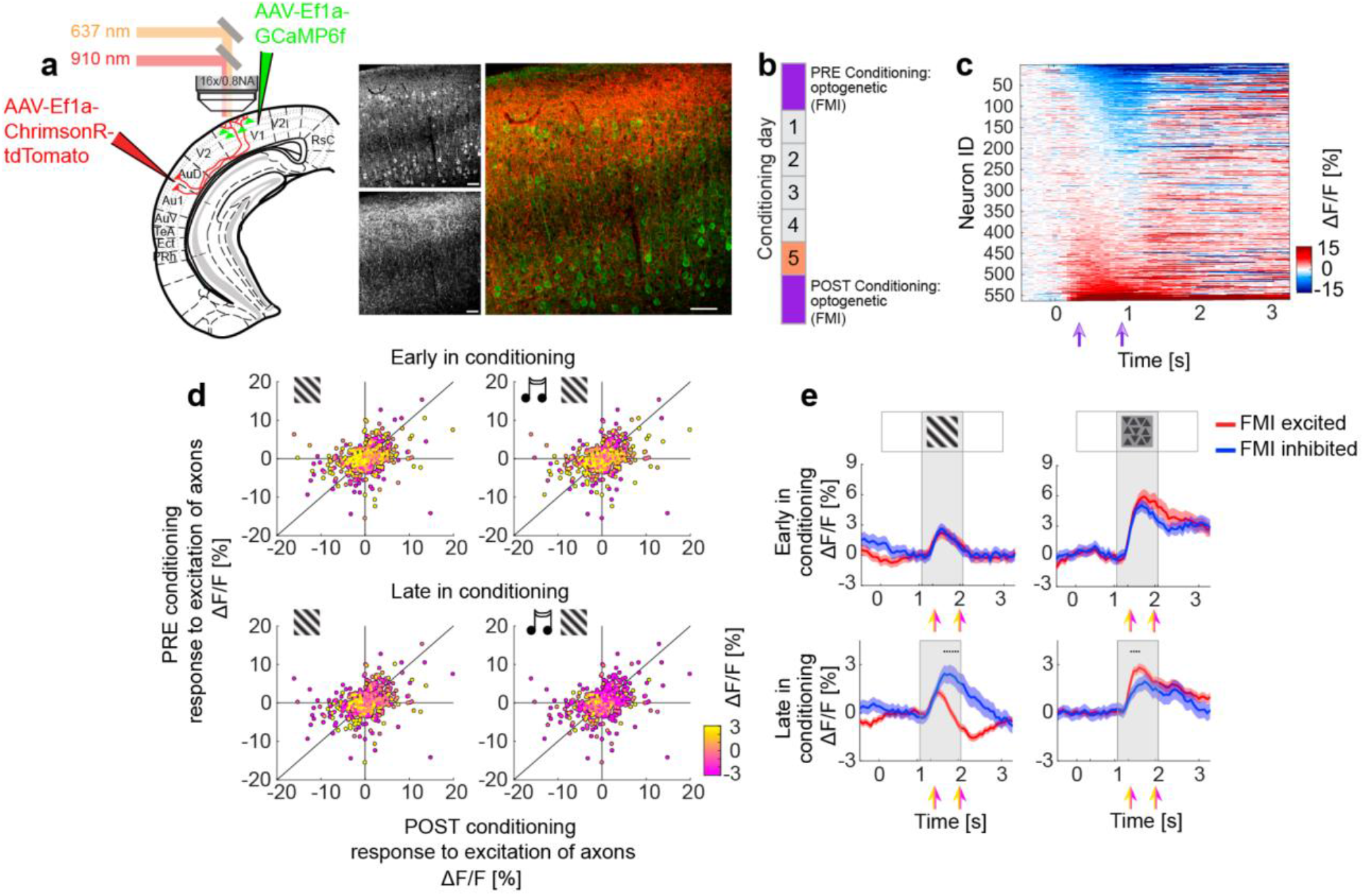
Auditory cortex input selectively inhibits visually responsive neurons in V1. **(a)** (Left) Schematic of injection sites referenced to atlas (Paxinos and Franklin, 2013). GCaMP6f injection in V1 and ChrimsonR-tdTomato injection in AuC. (Right) Representative confocal histology image illustrating AuC axons (bottom gray inset and red) and V1 neurons (top gray inset and green). Scale bars indicate 50 µm. **(b)** Optical stimulation of AuC projection axons in V1 was performed to functionally map the influence (FMI) of AuC input on V1 neurons one day before and one day after the 5 day conditioning paradigm. **(c)** V1 neuron responses to pre-conditioning optogenetic stimulation of AuC axons sorted by strength of response. Purple arrows indicate window over which response was averaged to generate FMI response values in **d**. For **c** and **d**, n = 563 neurons from 5 mice. **(d)** The response of each V1 neuron to optogenetic stimulation of AuC axons (FMI) before conditioning plotted against the response after conditioning. Color indicates the visual response of each neuron to V_a_ (left) or A_a_V_a_ (right), early (top) and late (bottom) in conditioning. **(e)** Visual responses of neurons inhibited (blue) or excited (red) by optogenetic excitation of AuC axons (FMI) to V_a_ (left) and V_c_ (right) early (top) and late (bottom) in conditioning. Colored arrows indicate window over which response was averaged for individual neurons to calculate visual response value plotted in **d**. Early in conditioning refers to first exposure to stimuli, which occurred on the pre-FMI day using visual stimulus trials without optogenetic stimulation. n = 563 neurons, 257 FMI inhibited, from 5 mice. Late in conditioning refers to an average of visual responses from days 3 and 4 of the conditioning paradigm (see also **Extended Data Figs. 4d, f**). n = 1548 neurons, 482 inhibited from 10 mice. Black dots indicate traces are different during visual stimulation (p<0.05, paired two-sided t-test, see Methods for detailed calculation). Traces represent the mean and shading represents SEM across neurons.

In summary, we find that the association of an auditory cue with a visual stimulus results in an experience-dependent suppression of the visual response in L2/3 V1 neurons that is specific to the paired association. While auditory modulation of visual cortex likely occurs via multiple pathways (McIntosh et al., 1998), one of the mechanisms that contributes to this experience-dependent suppression of predictable visual stimulation is direct input from auditory cortex. With experience, the functional influence of AuC input changes to selectively target the L2/3 V1 neurons responsive to the paired visual stimulus for inhibition. This inhibition is likely mediated by local inhibitory neurons that are recruited by AuC input (Deneux et al., 2019; Ibrahim et al., 2016; Iurilli et al., 2012). As the AuC input functions to suppress predictable visual input, these interactions are well described by the framework of predictive processing (Keller and Mrsic-Flogel, 2018). Similar long-range, cortico-cortical interactions are also thought to contribute to the suppression of predictable sound associated with movement (Schneider et al., 2014, 2018). Additionally, we found that a learned behavioral response to the auditory-cued visual stimulus was necessary for visual suppression, a result in concordance with previous work showing a correlation between experience-dependent changes in V1 responses and behavioral performance during appetitive learning but not passive viewing (Henschke et al., 2020). Our results are also consistent with the idea that cortical circuits are shaped by experience to store cross-modal associations, and thereby contribute to memory storage in sensory cortex (Buonomano and Merzenich, 1998; Gilbert et al., 2001; McGann, 2015). Moreover, blocking of the formation of an association of a stimulus with a reinforcement can occur when two conditioned stimuli are used as predictors (Mackintosh, 1975; Rescorla and Wagner, 1972). Because the auditory cue is predictive of reinforcements in our study, suppression of the visual response may be a mechanism of blocking. An associative memory trace is often considered to reside in higher association areas that receive convergent input from lower sensory areas. An alternative mechanism for such a trace is the synaptic change that combines or redirects information flow between long-range sensory projections and local sensory areas. We demonstrate that cross-modal learning can shape and redefine representational patterns of sensory stimuli through the interaction of long-range input with local circuits. Thus, long-range cross-modal interactions can shape representations of the sensory world endowing early sensory cortex with a mnemonic capacity (Murray et al., 2016; Weinberger, 2004) that functions to make cross-modal predictions.

## Acknowledgements

We thank Rainer W. Friedrich, Johannes Felsenberg, and Friedemann Zenke for helpful discussion and comments on earlier versions of this manuscript. We thank all the members of the Keller lab for discussion and support. This project has received funding from the Swiss National Science Foundation and the Novartis Research Foundation.

## Author Contributions

ARG designed and performed the experiments and analyzed the data. Both authors wrote the manuscript.

## METHODS

### Animals

All animal procedures were approved by and carried out in accordance with guidelines of the Veterinary Department of the Canton Basel-Stadt, Switzerland. Mice were both male and female and group housed by gender with a 12/12 h light/dark cycle in cages with horizontal running wheels.

### Surgeries

Surgeries were performed as described previously (Leinweber et al., 2014). Briefly, mice were anesthetized using a mix of fentanyl (0.05 mg/kg), medetomidine (0.5 mg/kg) and midazolam (5 mg/kg). A craniotomy of either 5 mm or 3 mm in diameter was made over V1, a glass coverslip was superglued in place, and a custom machined stainless-steel head bar was implanted.

### Adeno-associated virus (AAV) injections

Injections consisted of 100 to 250 nl of AAV vector with a titer in the range of 10^12^ to 10^14^ genome copies/ml. The coordinates of the injections in V1 were 2.7 to 2.8 mm lateral from the midline, and 2.8 to 3.0 mm posterior from bregma. For AuC injections the coordinates were 4.4 mm lateral from the midline and 2.6 to 2.8 mm posterior from bregma, and the injection pipette was rotated to be perpendicular to the brain surface. For somatic imaging in V1 we used AAV2/1-EF1α-GCaMP6f, for V1 PV-Cre excitation and functional mapping of influence we used AAV2/1-EF1α-ChrimsonR-tdTomato (Klapoetke et al., 2014), and for AuC axon imaging we used AAV2/1-EF1α-GCaMP6s (Chen et al., 2013).

### Histology

For post-mortem histological analyses, mice were transcardially perfused with 4% paraformaldehyde (PFA) in PBS. Brains were isolated and maintained in 4% PFA at 4°C on a shaker overnight. The fixed tissue was then rinsed with PBS and sectioned into 70 or 100 µm thick slices using a vibratome. Sections were mounted and sealed with DAPI ProLong mounting medium. Sections for all mice were imaged using a Zeiss AxioScan.Z1 slide scanner at 10x magnification. All images used for quantification of the number of neurons expressing GCaMP, were acquired at 20x magnification, 5 µm-step, z-stack images using a confocal microscope. Atlas overlays for histological images were adapted from Paxinos and Franklin 4^th^ ed. (Paxinos and Franklin, 2013). Atlas images were first aligned to both rhinal fissures and the external capsule of coronal sections and subsequently the thickness of the cortex was adjusted to fit each individual mouse. Confocal ex-vivo histology images were acquired for all mice.

### Quantification of AAV spread

Injections of AAV2/1-EF1α-GCaMP6s-WPRE in AuC for axonal imaging in V1 also result in axonal uptake and expression in V1 neurons that project to AuC. To quantify what fraction of the axons in V1 could come from retrogradely labelled V1 neurons, we used a separate set of 5 mice for histological quantification. Mice were injected with AAV2/1-EF1α-GCaMP6s-WPRE in AuC and sacrificed for histological analysis time matched to the start of the imaging experiments. We performed a histological quantification using confocal images of fixed tissue in a region corresponding to the location of our two-photon imaging window. We then quantified the number of neurons per slice volume (656 x 656 x 32 µm). We found infected neurons in V1 in two out of five mice with a mean ± SEM across mice of 2.6 ± 1.9 neurons, and 5 infected neurons in one out of five mice in secondary visual areas (1 ± 1, mean ± SEM across mice) (**Extended Data Figs. 2a, b**). Given that the number of axons we were able to image in V1 in a volume of 200 x 200 x 40 µm was more than two orders of magnitude larger (day 1: 1054.8 ± 117.8, day 2: 893.2 ± 91.6, day 3: 1008.2 ± 121.9, day 4: 1025.6 ± 130.0; mean ± SEM), retrogradely labelled V1 neurons are unlikely to account for a substantial fraction of the axons recorded in V1. Note, the comparison by volume is not entirely straightforward as one would need to estimate the average fraction of total V1 volume the axon of a given V1 neuron would be visible in. However, additionally arguing against a contamination by axons of V1 neurons is the fact that expression levels in retrogradely labelled neurons tend to be far lower than at the primary injection site (Tervo et al., 2016). Thus, although we cannot exclude that some of the axons in our dataset originated from retrogradely labelled V1 neurons, the vast majority of them were likely AuC projection axons.

### Two-photon imaging

Functional imaging of GCaMP6 expressing neurons was performed using a modified Thorlabs B-Scope. The illumination source for two-photon imaging was a femtosecond infrared laser (Spectra physics) tuned to a wavelength of 910 nm. A 12 kHz resonance scanner (Cambridge Technology) was used for line scanning to acquire data at a frame rate of 60 Hz at a resolution of 400 x 750 pixels. In addition, we used a piezo actuator (Physik Instrumente) to acquire images at 4 different depths by moving the objective (Nikon 16x, 0.8NA) in 15 µm steps between frames, thereby reducing the effective frame rate per imaging plane to 15 Hz.

### Optogenetic stimulation during two-photon imaging

The methods for simultaneous two-photon imaging and optogenetic stimulation were described previously (Attinger et al., 2017; Leinweber et al., 2017). Briefly, the illumination source for the ChrimsonR stimulation was a switchable 637 nm laser (OBIS, Coherent) used at an average power of 11 mW and triggered using a TTL pulse. A dichroic mirror (ZT775sp-2p, Chroma) was used to combine two-photon and optogenetic stimulation light, and a long-pass dichroic mirror (F38-555SG, Semrock) was used to filter GCaMP6 emission from illumination light. To prevent stimulation light artefacts, the 637 nm laser was synchronized to the turnaround times of the resonant scanner when data were not acquired. To reduce the influence of ringing artefacts in the amplifier, signals were digitally bandpass filtered at 80 MHz using a 1.6 GHz digitizer (NI-5772, National Instruments) and an FPGA (PXIe-7965, National Instruments) to implement a digital Fourier filter.

### Conditioning paradigm

#### Mice

Mice were handled by the experimenter every day for at least one week before being introduced to the VR. Water restriction began one week before the start of experiments in which a water reward was delivered, and mice received 1 ml of water per day. Three to five days before the experiment, mice were exposed and habituated to head fixation in the VR and rewarded with sunflower seeds after each exposure period. Mice were considered habituated when they voluntarily walked onto the experimenter’s hand and did not resist head-fixation. During experiments, mice received supplemental water after conditioning if they had not consumed at least 1 ml in water rewards. Mice were monitored to ensure they maintained at least 80% of their original body weight. For V1 soma imaging, one cohort of 5 mice underwent optogenetic experimentation in the VR context on day 1, followed by 5 days of conditioning, followed by a final day of optogenetics. A second cohort of 5 mice only had optogenetic experimentation after 5 days of conditioning. For AuC axon imaging, 20 mice were conditioned for 4 days. One mouse was removed from the analysis on day 4 due to insufficient image registration. Of these mice, 8 were PV-cre and were also used for optogenetic and visual context only experiments.

#### Stimuli

Auditory stimuli consisted of either a 16.1 or 10.5 kHz pure tones presented at approximately 65 dB SPL (Iurilli et al., 2012). The three visual stimuli used were a sinusoidal grating, a geometric pattern of triangles, and a geometric pattern of ovals.

One of the associated stimuli (a and b) was always the grating, but the pairing of the stimuli was otherwise randomized and counterbalanced across animals. For paired conditions, the auditory stimulus was 1 second in duration, followed immediately by a visual stimulus 1 second in duration, followed immediately by a reinforcement, a: water reward, b: airpuff. For visual stimulus only conditions, the visual stimulus was presented for 1 second and never reinforced. Approximately 25% of trials were the V_x_ condition during the first four conditioning days (day 1, V_a_: 24.5 ± 0.2%) and ∼14% of trials on day 5 (V_a_: 13.8 ±0.5%). The occurrence of V_c_ as a fraction of all un-cued visual stimulus trials was day 1: 50.1 ± 0.3 % and day 5: 33.9 ± 0.5%. On day 5, A_b_V_a_ occurred for ∼14% of all cued visual stimulus trials (A_b_V_a_: 13.8 ± 0.6,). Values reported are mean ± SEM. For axonal imaging, the visual only paradigm was performed one day before and after conditioning as well as following the audio-visual paradigm on conditioning days (**Fig. 2i**). Stimuli consisted of full field grating presentations of eight orientations with a stimulus duration of 2 seconds and a gray (mean-luminance) inter-stimulus-interval of 3 seconds.

#### Virtual reality (VR)

Mice were head-fixed and free to locomote on an air-supported polysterene ball. A virtual tunnel designed with low contrast gray checkered walls was projected onto a toroidal screen surrounding the mouse and yoked to linear displacement of the ball. From the mouse’s perspective, the screen encompassed a visual field of approximately 240°horizontally and 100° vertically. One speaker was placed on the left and one on the right side of the VR for presentation of auditory stimuli. The VR system was otherwise constructed as described previously (Leinweber et al., 2014). A water spout was placed in front of mice, and licking was detected using a custom made electrical circuit in which a mouse closes the circuit whenever her/his tongue contacts the metal spout or water droplet (Hayar et al., 2006). The resulting voltage was thresholded to calculate licking events.

#### Image analysis

Regions of interest (ROIs) for soma were obtained using custom semi-automated image-segmentation software. ROIs for axons were obtained in an automated process as previously described in Mukamel et. al. (Mukamel et al., 2009) using a combination of principle and independent component analysis and image segmentation modified in house. Fluorescence traces across time were then calculated as the mean pixel value in each ROI per frame. ΔF/F was calculated using median normalized traces and filtered as described previously (Dombeck et al., 2007). For axonal imaging, data came from the same location in the brain using blood vessel patterns for alignment, but individual axons were not matched across imaging time-points.

### Data analysis

Data analysis was performed using custom written Matlab (Mathworks) code. To quantify differences between response curves during visual stimulation (**Figs. 1c, g; 2c; 3e; Extended Data Figs. 2b, d, e; 4d, f**), ΔF/F was compared in a response time window (11 frames, 267 - 1000 ms post visual stimulus onset) with a baseline subtraction during auditory stimulation (mean activity in a window preceding visual stimulus onset, 10 frames, -667 - 0 ms) bin-by-bin for 1 frame (66.7 ms) time bins using a paired t-test (p<0.05). Dots above response curves indicate significant difference for at least 3 consecutive bins. For quantification of responses during visual, auditory, optogenetic, or sham stimulation, ΔF/F was averaged over the response time window (11 frames, 267 - 1000 ms post stimulus onset) and baseline subtracted (mean activity in a window preceding stimulus onset, 10 frames, -667 - 0 ms) (**Figs. 1d, h; 2g; 3d, e; Extended Data Figs. 1b - i; 2a, d, e; 3c - e; 4a, c, d - g**) Mean neural activity is an average across trials and neurons. Mean behavioral data is an average across trials and mice. Licking and running were quantified during the response time window (**Figs. 1f; Extended Data Figs. 2b, d, e**). For quantification of visually responsive axons (**Figs. 2g - i**), ΔF/F during the response time window was compared to ΔF/F during the baseline window. Normalized suppression of AuC axons was quantified as the difference between the response to the stimulus with and without optical stimulation of V1 PV neurons, normalized by the mean response to the stimulus without optical stimulation (**Fig. 2f**). The response difference index was computed by subtracting the response during the visual stimulus following the auditory cue (A_a,b,o_V_a,b,o_) from that during the visual stimulus presented alone (V_a,b,o_) (**Fig. 1d, h, Extended Data Figs. 1f, i, 2d, 4e**), the visual stimulus following the paired cue (A_a_V_a_) from that during the unpaired cue (A_b_V_a_) (**Fig. 1h, Extended Data Figs. 2e, 4g**), or the visual stimulus following the unpaired cue (A_b_V_a_) from that during the visual stimulus alone (V_a,b,o_) (**Fig. 1h**), and normalized to the mean visual response alone (V_a,b,o_) on day 1 of conditioning. Note, we used a subtractive measure normalized by day 1 responses to avoid division by 0 problems. For classification of V1 neurons as excited by or inhibited by AuC stimulation, we split the population of neurons into two groups. Those with a response greater than 0 were included in the excited-by group and those with a response less than 0 in the inhibited-by group (**Fig 3e; Extended data Figs. 4d - g**). For locomotion speed-matching (**Extended Data Figs. 2d, e**), an iterative resampling procedure was used: the fastest and slowest trials were successively removed in the stimulus conditions with higher and lower average running speeds, respectively, until average running speed in the condition with the initially higher average running speed was lower than in the condition with the initially lower average running speed. For **Fig. 3d, e**, early in conditioning is day 1 of experiment (first exposure to conditioning stimuli), and late in conditioning is the average of the visual responses on days 3 and 4 of conditioning. For the no reinforcement paradigm (**Extended Data Figs. 1g - i**), mice were exposed to two sets of stimuli as in the reinforced experiments, A_a_V_a_ and A_b_V_b_, but as neither condition was reinforced, visual and auditory cue responses were calculated by averaging across both conditions (A_o_V_o_ is the average of A_a_V_a_ and A_b_V_b_, V_o_ of V_a_ and V_b_, and A_o_ of A_a_ and A_b_).

### Statistical analyses

All statistical analyses were performed in Matlab (Mathworks) using custom written software. Sample sizes were chosen to match typical numbers used in animal behavioral experiments. All data acquired were included in the analysis. Changes in the number of mice (and neurons) across time-points are the result of technical difficulties that prevented the acquisition of data in some mice (**Table 1**). Data were first tested for normality using a Lilliefors test, and when the null hypothesis could not be rejected (h_o_: data come from a normally distributed population), parametric tests were used. Otherwise non-parametric tests were used. Paired *t*-tests or rank-sum tests were used for analyses with matched samples. For all unmatched-samples data that failed to reject the h_o_ in the Lilliefors test, unpaired t-tests were used (for example, comparisons of axon responses on different conditioning days). Error shading and bars indicate standard error to the mean (SEM) unless otherwise stated in figure legends. All statistical tests were two-tailed. Scattered data were quantified using correlation coefficients, denoted as r, and coefficients of determination were computed by taking the square of r.

**Table 1.** Statistical tests used, n’s and p-values.

| tp | p | # soma | # mice |
| --- | --- | --- | --- |
| 1 | 0.258 | 1548 | 10 |
| 2 | 0.183 | 1548 | 10 |
| 3 | 1.19E-06 | 1548 | 10 |
| 4 | 4.77E-28 | 1548 | 10 |
| 5 | 4.93E-15 | 1341 | 9 |

**f** test: rank-sum, # lick events during AaVa compared to Va across mice
| tp | p | # mice |
| --- | --- | --- |
| 1 | 0.4263 | 10 |
| 2 | 0.3075 | 10 |
| 3 | 0.064 | 10 |
| 4 | 0.0452 | 10 |
| 5 | 0.0039 | 9 |

**h** test: rank-sum, response difference index compared to 0 difference across soma
| condition | p | # soma | # mice |
| --- | --- | --- | --- |
| AbVb to AaVa | 1.49E-16 | 1341 | 9 |
| AbVa to Va | 0.372 | 1341 | 9 |

| condition | p | # axons | # mice |
| --- | --- | --- | --- |
| pre | 0.1784 | 2927 | 7 |
| post | 1.58E-20 | 3857 | 7 |
| tone | 2.42E-176 | 4130 | 6 |

**h** test: paired t-test, # visually responsive axons to AaVa compared to Va
| tp | p | # axons | # mice |
| --- | --- | --- | --- |
| 1 | 0.133 | 5552 | 8 |
| 2 | 0.0255 | 5097 | 8 |
| 3 | 0.0214 | 5157 | 8 |
| 4 | 0.0135 | 4658 | 7 |

**i** test: unpaired t-test, # visually responsive axons on day 1 compared to day 4
| plot | p |
| --- | --- |
| aud-vis | 0.0202 |
| vis only | 0.5361 |

| tp | # axons | # mice |
| --- | --- | --- |
| 1 | 5552 | 8 |
| 4 | 4658 | 7 |

**EXTENDED DATA:****Figure 1** **f** test: rank-sum, response difference index compared to 0 difference across soma
| tp | p | soma | mice |
| --- | --- | --- | --- |
| 1 | 3.58E-23 | 1548 | 10 |
| 2 | 2.42E-31 | 1548 | 10 |
| 3 | 7.04E-08 | 1548 | 10 |
| 4 | 4.74E-09 | 1548 | 10 |
| 5 | 1.57E-15 | 1341 | 9 |

| tp | p |
| --- | --- |
| 3 |  |
| i (light) | 0.0962 |
| ii (med) | 0.0027 |
| iii (drk) | 1.67E-08 |

**i** test: rank-sum, response difference compared to 0 difference across soma
| tp | p | soma | mice |
| --- | --- | --- | --- |
| 1 | 0.061 | 496 | 7 |
| 2 | 0.0069 | 496 | 7 |
| 3 | 0.0457 | 496 | 7 |
| 4 | 0.0015 | 496 | 7 |
| 5 | 7.81E-04 | 335 | 5 |

| p | tp | soma | mice |
| --- | --- | --- | --- |
| 0.0391 | 1 | 1548 | 10 |
|  | 4 | 1548 | 10 |

**d & e** **rank-sum**, response difference index compared to 0 difference across soma
| tp | p | soma | mice |
| --- | --- | --- | --- |
| 4 | 5.41E-10 | 1548 | 10 |
| 5 | 5.44E-14 | 1341 | 9 |

| p | r | soma | mice |
| --- | --- | --- | --- |
| 0.71 | -0.016 | 563 | 5 |

**c** **test: correlation coefficients**, null: no relationship exists between variables
| p | r | soma | mice |
| --- | --- | --- | --- |
| 6.51E-252 | 0.9334 | 563 | 5 |

**e** **test: rank-sum**, comparison of response difference indices between excited and inhibited soma
| tp | p | soma | # excited | # inhibited | mice |
| --- | --- | --- | --- | --- | --- |
| 5 | 1.78E-14 | 1341 | 927 | 414 | 9 |

**g** **test: rank-sum**, comparison of response difference indices between excited and inhibited soma
| tp | p | soma | # excited | # inhibited | mice |
| --- | --- | --- | --- | --- | --- |
| 5 | 6.48E-10 | 1341 | 927 | 414 | 9 |

### Data availability

All raw data necessary to reproduce all figures is available here: https://data.fmi.ch/.

### Code availability

All analyses code necessary to reproduce all figures is available here: https://data.fmi.ch/. Core analysis and imaging code is available here: https://sourceforge.net/projects/iris-scanning/

## EXTENDED DATA

**Extended Data Figure 1.**
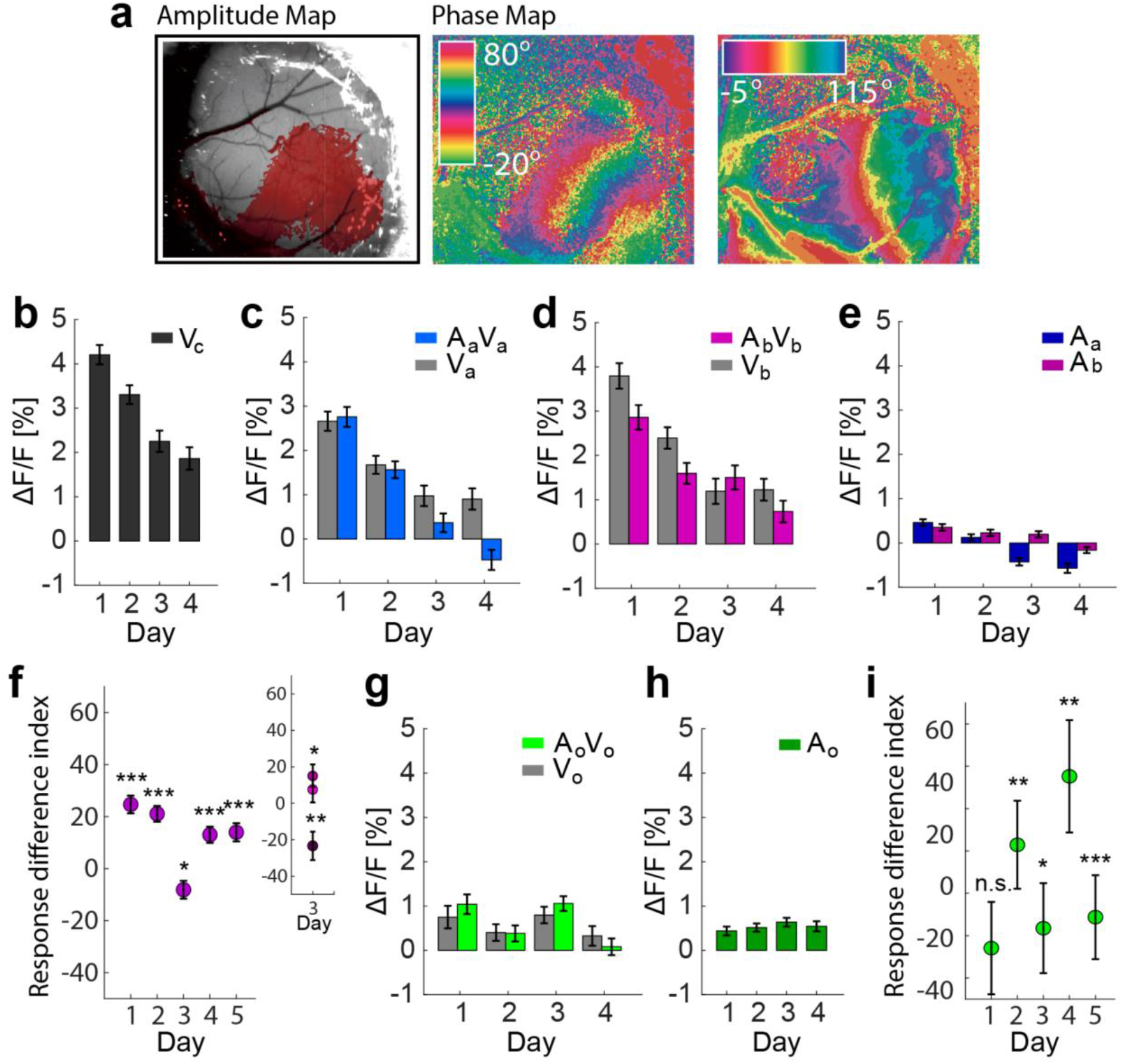
**(a)** Intrinsic signal optical imaging was performed on all mice before 2-photon imaging, n = 30 mice. Shown are data from one representative mouse. **(b)** Average population visual responses as a function of conditioning day to V_c_ (never paired), **(c)**visual responses to A_a_V_a_ (positive reinforcement, blue) and V_a_ (gray), **(d)**visual responses to A_b_V_b_ (negative reinforcement, pink) and V_b_ (gray), and **(e)**responses to the auditory cue, A_a_ (blue) and A_b_ (maroon). For **b - e** n = 1548 neurons from 10 mice. **(f)**Quantification of the difference in response for each conditioning day (Response difference index) during the auditory-cued and un-cued visual stimulus presentations, normalized by the mean response during the un-cued visual stimulus on day 1 (V_a_-A_a_V_a_)/ mean(V_a_). On day 3, the visual response to A_b_V_b_ was on average larger than that to V_b_ (see also panel **d**) resulting in a negative suppression. However, this effect was driven by a few outliers, which can be seen when the data is split into three epochs (inset). The negative suppression is only present in the 3^rd^ epoch of the day. Day 1 - 4: n = 1548 neurons from 10 mice; day 5: n = 1341 neurons from 9 mice. Asterisks indicate comparison to 0 difference using a two-sided rank-sum test. Here and in subsequent panels *: p < 0.05, **: p < 0.01, ***: p < 0.001; for all statistical analyses and exact p values see **Table 1**. **(g)**Average population visual responses as a function of conditioning day when stimuli were not reinforced. A_o_V_o_ (no reinforcement, green) and V_o_ (gray), and **(h)**responses to the auditory cue A_o_ (dark green). For **g** and **h**, n = 496 neurons from 7 mice. Subscript o indicates an average across conditions a and b (i.e. A_a_V_a_ and A_b_ V_b_, V_a_ and V_b_, A_a_ and A_b_) because neither condition a or b was reinforced. **(i)**Quantification of the difference in response for each conditioning day (Response difference index) during the auditory-cued and un-cued visual stimulus presentations in the no reinforcement paradigm. Calculated as in panel **f**. Day 1 - 4: n = 496 neurons from 7 mice, day 5: n = 335 neurons from 5 mice. Error bars indicate SEM over neurons.

**Extended Data Figure 2.**
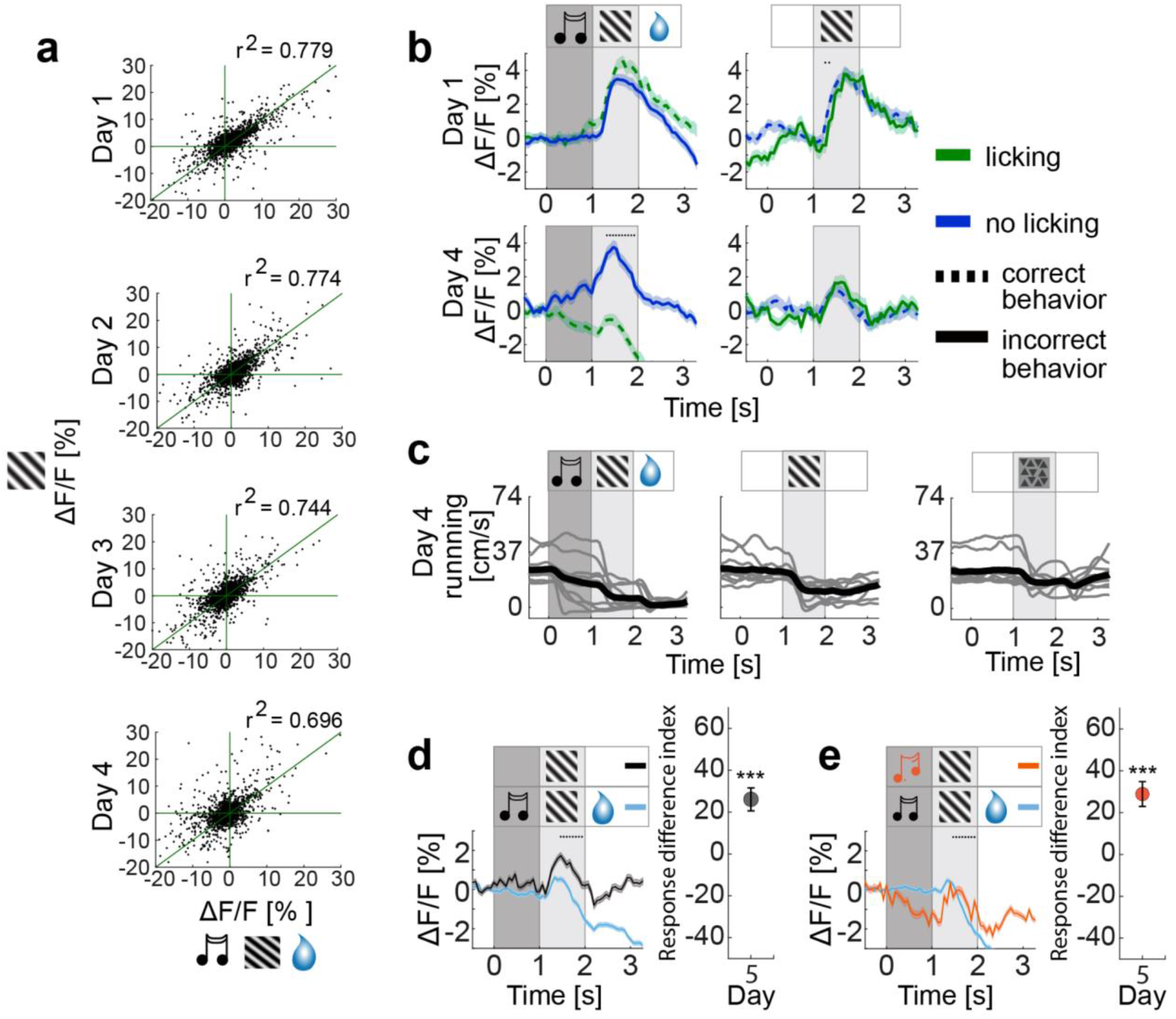
(**a**) The proportion of explained variance comparing responses during the visual stimulus presented alone, V_a_, and presented following the auditory cue, A_a_V_a_, decreases with conditioning day. r^2^ for the entire population of neurons is indicated on scatter plots; r^2^ per mouse mean ± SEM: day 1: 0.738 ± 0.051, day 4: 0.517 ± 0.091, p < 0.05 paired t-test, r values day 1 vs. day 4 comparison, n= 1548 neurons from 10 mice for **a, b**, and **d**. (**b**). Average population responses to A_a_V_a_ (left) and V_a_ (right) on day 1 (top) and day 4 (bottom) for trials during which mice licked (green) and failed to lick (blue). Dashed lines indicate correct licking preceding reward or correct withholding of licking preceding no reward during stimulus presentations. Solid lines indicate the converse (incorrect) licking behavior. **(c)** Average running speeds during stimulus presentations (gray, each mouse; black, mean across mice), n = 10 mice, running speed before stimulus onset, 25.2 ± 2.9 cm/s and during visual stimulation, A_a_V_a_: 7.6 ± 2.4 cm/s, V_a_: 12.7 ± 1.6 cm/s, and V_c_: 18.1 ± 1.4 cm/s (mean ± SEM). **(d)** (Left) Average population responses of L2/3 V1 neurons for cued (A_a_V_a_, blue) and un-cued (V_a_, gray) visual stimulus presentations on day 4 of conditioning for running speed matched trials. Average speed and total number of trials included for A_a_V_a_: 9.0 cm/s, 514 trials and for V_a_: 8.9 cm/s, 134 trials. (Right) Response difference index. Asterisk indicates comparison to 0 difference using a two-sided rank-sum test. n = 1548 neurons from 10 mice. **(e)** (Left) Average population responses of L2/3 V1 neurons for the previously paired cue (A_a_V_a_, blue) and previously un-paired cue (A_b_V_a_, orange) visual stimulus conditions on day 5 of conditioning for running speed matched trials. Average speed and total number of trials included for A_a_V_a_: 11.3 cm/s, 960 trials and for A_b_V_a_: 10.6 cm/s, 123 trials. (Right) Response difference index. Asterisk indicates comparison to 0 difference using a two-sided rank-sum test. n = 1341 neurons from 9 mice. Traces or filled circles indicate the mean and shading or error bars indicate SEM across neurons.

**Extended Data Figure 3.**
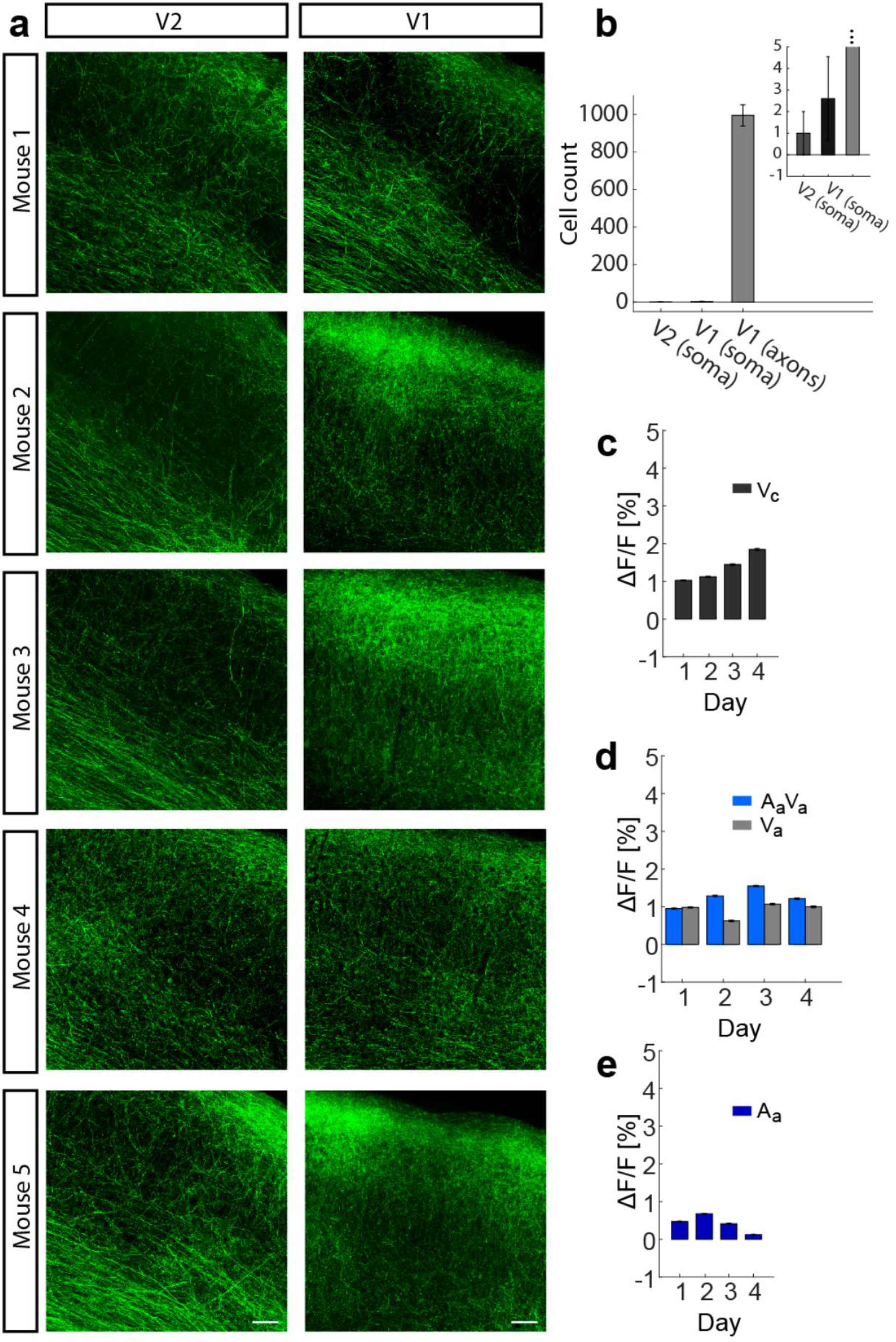
**(a)** (Injection in AuC (see methods) to label projection axons (green). Z projection of confocal images shows approximately 656 x 656 x 32 um of secondary visual cortex (V2)(left) and V1 (right). Scale bar indicates 50 µm. **(b)** Quantification of infected soma in V2 (left) and V1 (middle), and axons in V1 (right) after injection in AuC. Inset: same but scaled to range of soma numbers. Mean ± SEM across mice. **(c)** Average population visual responses as a function of conditioning day to V_c_(never paired), **(d)** visual responses to A_a_V_a_ (positive reinforcement, blue) and V_a_ (gray), **(e)** and responses to the auditory cue, A_a_. For **c - e**, day 1: n = 5552 axons from 8 mice, day 2: n = 5097 axons from 8 mice, day 3: n = 5157 axons from 8 mice, and day 4: n = 4658 axons from 7 mice.

**Extended Data Figure 4.**
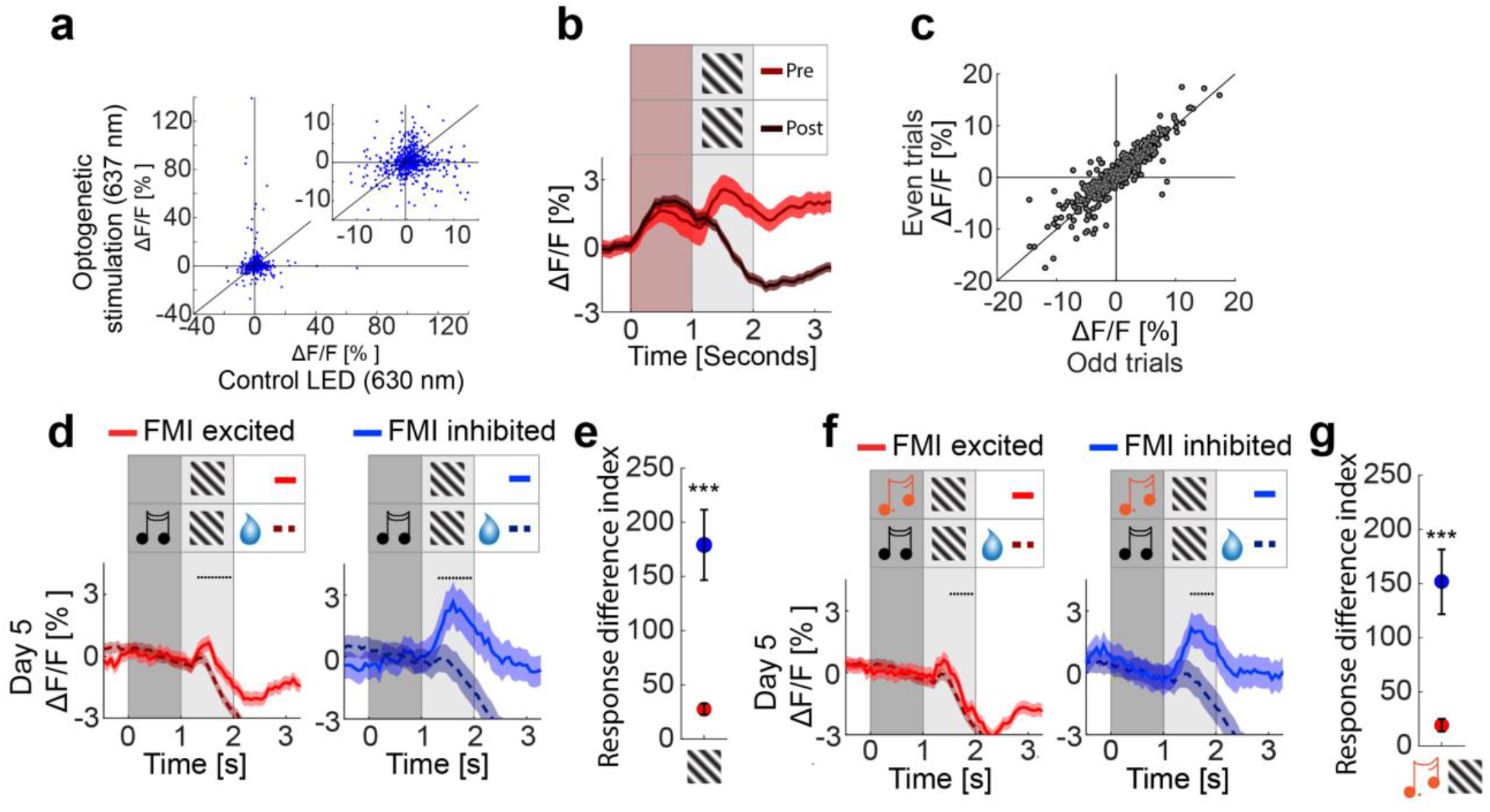
**(a)** Population responses to optogenetic stimulation (y-axis) compared to sham stimulation (x-axis) show no correlation. r = -0.016, p = 0.71. For **a** and **c**, n = 563 neurons from 5 mice. **(b)** The average population response of V1 soma to optogenetic stimulation of AuC axons pre-(light red) and post-(dark red) conditioning, Pre: n = 563 neurons from 5 mice, Post n = 1548 neurons from 10 mice. **(c)** The response of all V1 neurons to optogenetic stimulation of AuC axons on even numbered trials plotted against the response on odd numbered trials. r = 0.933, p <0.001. **(d)** Average population responses of V1 neurons excited (reds, left) and inhibited (blues, right) by AuC stimulation to V_a_ (solid trace) and A_a_V_a_ (dashed trace) presentations on conditioning day 5. For **d, e, f**, and **g**: n = 1341 total, 927 excited, and 414 inhibited neurons from 9 mice. **(e)** Response difference index for data shown in **d**. Comparison between excited and inhibited neurons using a rank-sum test. Here and in subsequent panels *: p < 0.05, **: p < 0.01, ***: p < 0.001; for all statistical analyses and exact p values see **Table 1**. **(f)** Average population responses of V1 neurons excited (reds, left) and inhibited (blues, right) by AuC stimulation to A_b_V_a_ (solid trace) and A_a_V_a_(dashed trace) presentations on conditioning day 5. **(g)** Response difference index for data shown in **f**. Comparison between excited and inhibited neurons using a rank-sum test.

